# Field-crop transcriptome models are enhanced by measurements in systematically controlled environments

**DOI:** 10.1101/2024.09.21.614268

**Authors:** Yoichi Hashida, Daisuke Kyogoku, Suguru E. Tanaka, Naoya Mori, Takanari Tanabata, Hiroyuki Watanabe, Atsushi J. Nagano

**Affiliations:** Faculty of Agriculture, Takasaki University of Health and Welfare, Takasaki, Gunma, Japan; Museum of Nature and Human Activities, Sanda, Hyogo, Japan; Research Institute for Food and Agriculture, Ryukoku University, Otsu, Shiga, Japan; Biosystems & Biofunctions Research Center, Research Institute, Tamagawa University, Machida, Tokyo, Japan; Facility for Genome Informatics, Kazusa DNA Research Institute, Kisarazu, Chiba, Japan; Department of Advanced Food Sciences, Faculty of Agriculture, Tamagawa University, Machida, Tokyo, Japan; Faculty of Agriculture, Ryukoku University, Otsu, Shiga, Japan; Institute for Advanced Biosciences, Keio University, Tsuruoka, Yamagata, Japan

**Keywords:** Rice, Field, Growth chamber, RNA-Seq, Statistical modelling, Transcriptome

## Abstract

Plants in the field respond to seasonal and diel changes in various environmental factors such as irradiance and temperature. We previously developed a statistical model that predicts rice gene expression from the meteorological data and identified the environmental factors regulating each gene. However, since irradiance and temperature (the two most critical environmental factors) are correlated in the field, it remains difficult to distinguish their roles in gene expression regulation. Here, we show that transcriptome dynamics in the field are predominantly regulated by irradiance, by the modelling involving diurnal transcriptome data from the 73 controlled conditions where irradiance and temperature were independently varied. The model’s prediction performance is substantially high when trained using field and controlled conditions data. Our results highlight the utility of a systematic sampling approach under controlled environments to understand the mechanism of plant environmental response and to improve transcriptome prediction under field environments.

## Introduction

Plants in nature and crops in agricultural fields adapt to seasonal and diel changes in various environmental factors such as irradiance and temperature. Since these environmental factors fluctuate at fine spatiotemporal scales, plants must extract essential information from the field environment for their survival and respond to them accordingly. Understanding how plants sense and respond to these environmental variations is crucial for elucidating their adaptation mechanisms and increasing crop production^1^.

Although controlled environments such as glasshouses or growth chambers (GC) are commonly used in plant environmental response studies, findings obtained in such studies have not fully captured the complexities of plant responses to field conditions^2^. To address this issue, omics analysis has emerged as a promising way to study plant environmental responses to the field^3–5^. In particular, seasonal and diel transcriptome analysis has proven effective in elucidating how plants respond to different environmental stimuli in the field^6–16^.

The complementary approach is to mimic fluctuating field conditions in controlled environments. This approach has the advantages of repeatability and varying only environmental factors in interest^17–21^. Such studies have provided insight into several aspects of plant physiology, including photosynthesis^22–24^, flowering^25–27^ and metabolic regulation^28,29^.

Statistical modelling approaches have effectively extracted meaningful insights from noisy transcriptome data collected in the field^6,8–10^. We previously developed a statistical model that predicts transcriptome dynamics of rice leaves in the field from meteorological data^6,8,9^. The model suggested that transcriptome dynamics are predominantly governed by endogenous diurnal rhythms, ambient temperature, solar radiation, and plant age. The model well predicted the transcriptome dynamics of rice leaves grown in different years from the data used to develop the model. However, the model has low prediction accuracy at rare conditions in the field, such as extraordinarily high temperatures. This is due to the narrow range of environmental parameters and their correlation in the field (e.g., high temperature during the day and low temperature during the night). One solution to this problem is adding transcriptome data from controlled conditions with a wide range of environmental parameters while ensuring that the parameters are not correlated. However, setting up many growth chambers is logistically difficult^19^. In this study, we developed cost and space-effective GCs. Using the GCs, we conducted a massive transcriptome analysis of rice leaves grown under 73 conditions: three photoperiods, five temperature levels in the light, and five temperature levels in the dark. Using transcriptome data of the GC-grown rice and field-grown rice as input data, we extended our statistical model for predicting gene expression from meteorological data, highlighting the importance of irradiance in transcriptome regulation, a contrary conclusion from the previous study^9^. Our results contribute to a deeper understanding of plant responses to the environment in the natural environment and agricultural field.

## Results

### Systematic measurement of rice transcriptome dynamics across 73 growth chamber conditions

We conducted a massive transcriptome analysis of rice leaves grown in 73 conditions, combining photoperiod length and temperatures in light and dark periods. We developed a cost- and space-effective growth chamber (GC), enabling multi-environment tests in controlled conditions (Supplementary Fig. 1). By utilizing the GC, we grew two cultivars of rice (Koshihikari and Takanari). Koshihikari is a leading japonica cultivar in Japan, while Takanari is a high-yield indica cultivar with some indicia-specific features, such as a high photosynthetic rate^30^. Before the experiment, we grew all the rice plants for 14 days at 25/21°C under 12L/12D photoperiod (Supplementary Fig. 1). Subsequently, we transferred the plants to the GCs with different conditions, where they were left to acclimate for two days. The leaves were sampled on the third day. In each condition, we sampled rice leaves from one plant per cultivar every 3 h for 8 times (1168 total samples: 8 time-points × 2 cultivars × 73 conditions) and conducted RNA-Seq analysis (Supplementary Table 2). During the RNA-Seq analysis, replacement of the labelling of samples occurred. By predicting air temperature at the sampling timepoint from transcriptome data, we corrected the labelling of samples (Supplementary Figs. 2 and 3). After omitting samples with different genotypes from the labelling of the sample (Supplementary Fig. 2), 1167 samples were used for further analysis (Fig. 1a,b).

**Fig. 1.**
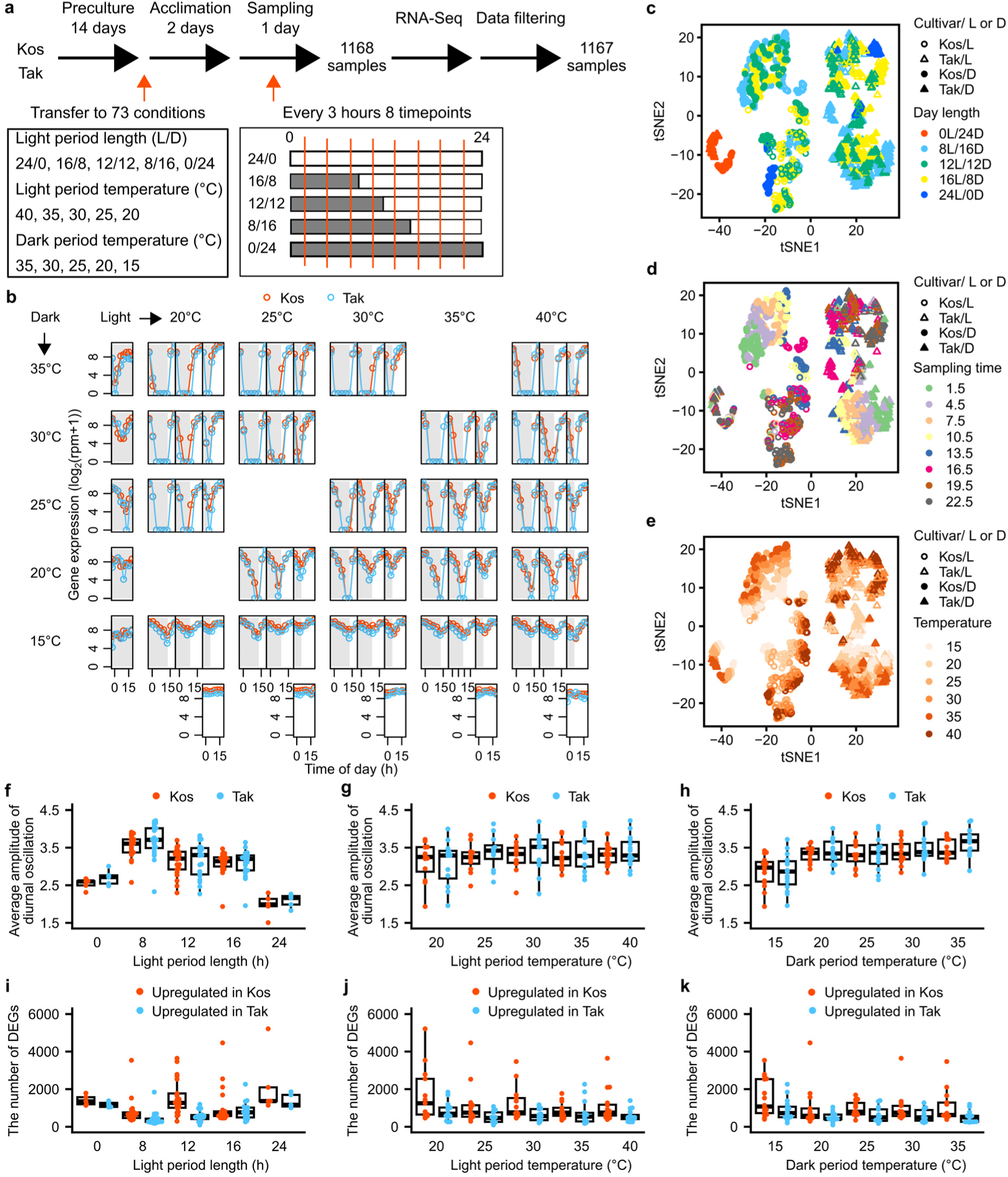
Systematic measurement of rice transcriptome dynamics across 73 growth chamber conditions for plant environmental response insights. a, Experimental design of this study. b, Expression levels of *OsGI* (*Os01g0182600*) in 73 conditions. c–e, t-Distributed Stochastic Neighbor Embedding (t-SNE) visualization showing clusters of transcriptomes of each sample coloured by (c) light period length, (d) light period temperature, and (e) dark period temperature. f–h, Genome-wide average amplitudes of diurnal oscillation of gene expressions at each condition, which is classified by (f) light period length, (g) light period temperature, and (h) dark period temperature. i–k, The number of differentially expressed genes (DEGs) between Koshihikari and Takanari at each condition, which is classified by (i) light period length, (j) light period temperature and (k) dark period temperature.

### Independent effects of light period length and temperature on rice gene expression and cultivar-specific responses

Our massive transcriptome data under controlled conditions allow us to disentangle the independent effect of light period length and temperature on rice gene expression. We evaluated transcriptome similarity between conditions using t-SNE (Fig. 1c**–**e).

Transcriptomes in continuous dark conditions (0L/24D) were separated from the other conditions. The other samples were mainly classified by cultivar and light/dark at the sampling time. We calculate the amplitude of diurnal oscillation of gene expression, which was lower in the continuous dark (0L/24D) and light (24L/0D) conditions than in the other conditions (Fig. 1f). The amplitude of diurnal oscillation was higher in 8L/16D than in 12L/12D and 16L/8D; the amplitude was higher in short-day conditions. The amplitude of diurnal oscillation was neither different between cultivars nor affected by light period temperature. However, it was affected by dark period temperature, where the amplitude of diurnal oscillation was lower at 15°C than in the other conditions (Fig. 1g,h). This is consistent with the previous study, which showed that the amplitude of diurnal oscillation of gene expression was lower in winter than in summer in *Arabidopsis halleri* subsp. *gemmifera*^7^. Further investigation is required to clarify whether 15°C during the daytime also decreases the amplitude of the diurnal oscillation of gene expression.

To clarify the difference in the environmental response of Koshihikari and Takanari, we compared the number of differentially expressed genes (DEGs) between the two cultivars at each condition. Because Koshihikari and Takanari belong to different subspecies (japonica and indica, respectively), some genes showed cultivar-specific gene expression^6,31–33^. To elucidate the effect of environmental conditions, we excluded cultivar-specific genes from the DEGs (Fig. 1i**–**k). Cultivar-specific genes were defined as those where the between-cultivar difference in the average expression (log_2_(rpm+1)) calculated across the 73 conditions was greater than 2, and the average expression in the cultivar with lower expression was less than 0.5. The number of cultivar-specific genes in Koshihikari and Takanari were 483 and 110, respectively. The number of DEGs tended to be higher in 0L/24D and 24L/0D than in the other conditions. The absence of environmental fluctuations under constant conditions may exaggerate cultivar differences. The expression level in Koshihikari tended to be higher than in Takanari in 12L/12D conditions, but the difference between the cultivars was not seen in the other conditions (Fig. 1i). The number of DEGs tended to be higher in Koshihikari than in Takanari in the low-temperature conditions: 20 °C in the light period and 15 °C in the dark period. This might result from the cold tolerance of the japonica cultivar compared with indica cultivars^33,34^. Collectively, our results showed the effect of light period length on the amplitude of gene expression and different responses to a low temperature between Koshihikari and Takanari.

### Co-expression gene network analysis identifies genes responsive to temperature

We analyzed the co-expression gene network using WGCNA^35^. The co-expression gene network was composed of 22 modules (clusters of co-expressed genes) harbouring 20–1416 genes (Fig. 2a). To find modules that have different expression patterns between Koshihikari and Takanari, we compared the mean value of eigengenes between the two cultivars, which was defined as the first principal component of the genes within a module (Fig. 2b). Module 1 was preferentially expressed in Koshihikari. In module 1, genes annotated for defence response (GO:0006952) and tetrapyrrole binding (GO:0046906) were significantly enriched (Fig. 2a and Supplementary Table 3). This is consistent with the previous studies showing the preferential expression of genes related to disease resistance^32,33^ and porphyrin and chlorophyll metabolism^33^ in japonica rice.

**Fig. 2.**
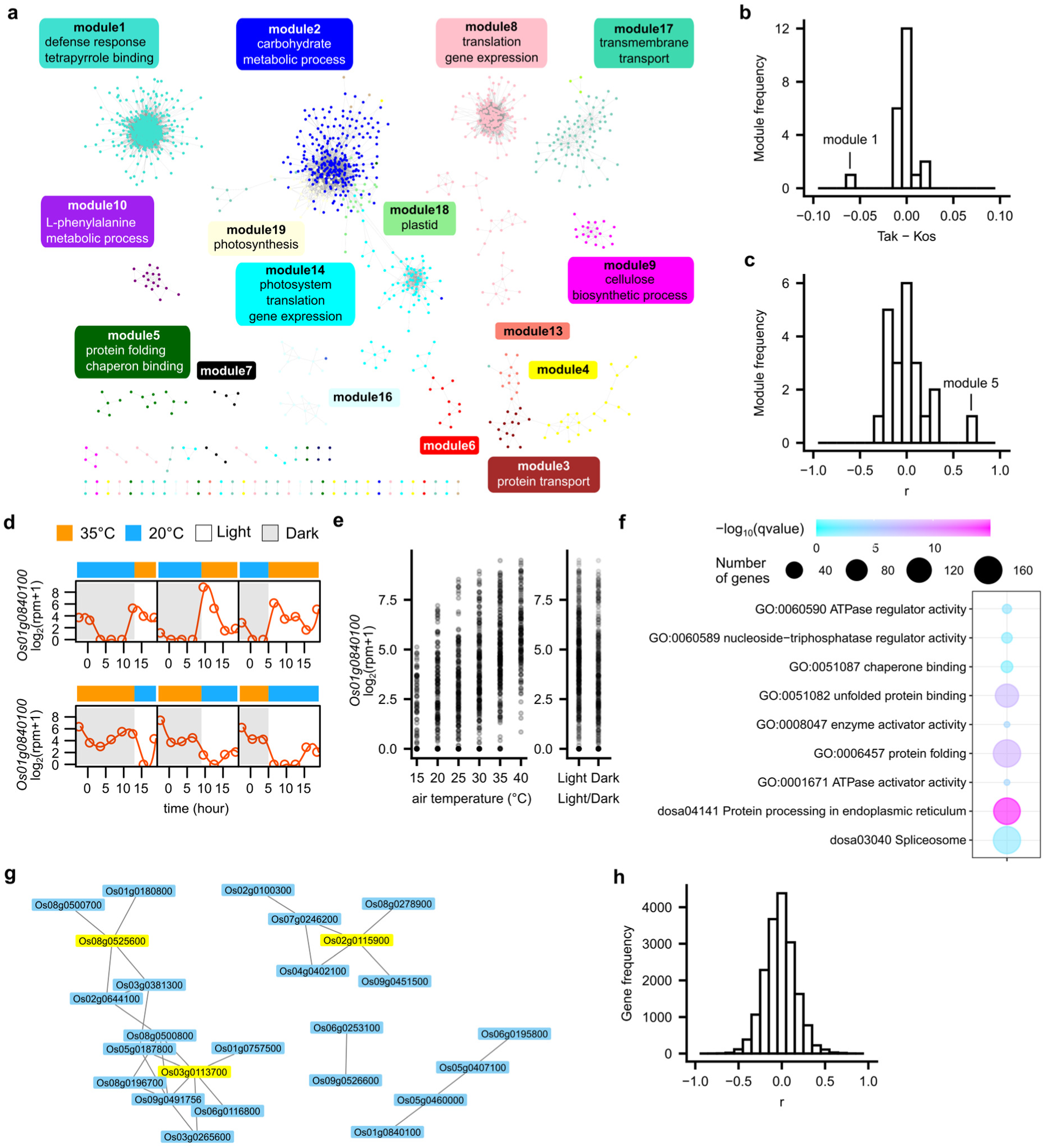
Identification of genes responsive to environmental conditions. **a**, Co-expression network constructed by WGCNA. Representative Gene Ontology (GO) enriched in each module is shown. **b**, A histogram showing the differences in mean expression of module eigengenes of Koshihikari and Takanari. **c**, A histogram showing Pearson’s correlation coefficient between the expression of module eigengenes and air temperature. **d**,**e**, Expression levels of a gene encoding *Hsp 70* (*Os01g0840100*) in module 5 (**d**) at 35/20°C (L/D) and 20/35°C (L/D) conditions in Koshihikari and (**e**) 73 conditions in Koshihikari and Takanari at each air temperature or light/dark condition. **f**, Gene Ontology and KEGG pathway significantly enriched in the temperature-responsive module 5. **g**, Gene network and hub genes of the temperature-responsive module 5. **h**, A histogram showing Pearson’s correlation coefficient between the expression of each gene and air temperature.

To detect specifically temperature-responsive modules, we calculated Pearson’s correlation coefficient (*r*) between temperature and the expression level of the eigengenes (Fig. 2c). Module 5 showed a strong positive correlation with temperature (*r* = 0.69), suggesting that module 5 is composed of temperature-responsive genes (Fig. 2c**–**e; Supplementary Figs. 4 and 5). On the other hand, the other modules were not correlated with temperature. The expression of the genes belonging to the modules tended to be constant (e.g., module 2) or regulated by the light/dark cycle (e.g., modules 4, 8, 17, and 18) (Supplementary Fig. 4).

To characterize the modules, we tested the enrichment of genes with gene ontology (GO) annotation and KEGG pathway in the modules. Genes annotated for chaperone binding (GO:0051087), protein folding (GO:0006457), and protein processing in the endoplasmic reticulum (KEGG pathway: dosa04141) were significantly enriched in module 5 (Fig. 2a,f; Supplementary Tables 3 and 4), indicating that genes belonging to module 5 is composed of heat stress-related genes (Supplementary Table 5). We identified three hub genes in module 5 that can have a central role in the regulation of the module (Fig. 2g). *Os02g0115900* encodes binding immunoglobulin protein (BiP), which is an endoplasmic reticulum-localized Hsp70 chaperone^36^. *Os03g0113700* encodes mitochondrial Hsp70^37^. *Os08g0525600* encodes FK506 binding protein, which is involved in stress response^38^. We also calculated the Pearson’s correlation coefficient (*r*) of gene expression and temperature to extract genes responsive to air temperature. *r* ranged from −0.76 to 0.77 (Fig. 2h; Supplementary Tables 6 and 7). Among the 91 genes with positive correlation (*r* > 0.5), 38 genes were members of module 5, and these genes include heat shock protein genes (Supplementary Tables 5 and 6). Expressions of 60 genes were negatively correlated with air temperature (*r* < −0.5) (Supplementary Table 7).

These genes might be important for adjustment to air temperature change in rice. Collectively, our results identified the temperature-responsive genes and the difference in gene expression between rice cultivars. Further analysis of temperature-responsive genes will contribute to plant response to temperature.

### Radiation is a better predictor of expression than temperature for the majority of genes

We then compared radiation and temperature as the predictors of gene expression dynamics. Under the field conditions, radiation and temperature are correlated with each other. Here we are interested in how the data from the GC, where radiation and temperature were systematically varied independently, enabled the choice of the better predictor. We thus trained statistical models for gene expression dynamics using different training data sets: GC data, field data, or a mixture of them (50% GC and 50% field). We mainly report the results for Koshihikari in the main text, but the results for Takanari were qualitatively the same (Supplementary Figs. 6 and 7). We focused on 474 genes representing various expression patterns in the field. These genes were chosen based on the microarray data analyzed by Nagano et al.^9^ and were representative of each cluster of genes with similar expression (Online Methods). Among the 474 representative genes, 8 genes showed too low expression levels for model training. We thus trained models for the remaining 466 genes (one model for a gene). We used the R package FIT^8^ for the model training, which chooses either radiation or temperature (the two most influential environmental variables to gene expression^9^) as the predictor of the gene expression (using neither is also an option). We randomly subsampled 512 data points from each set to standardize the size of different training data. We repeated the subsampling and model training 100 times. We can thus calculate how frequently radiation or temperature was chosen as the predictor for each gene (Fig. 3). When trained with field data, temperature was more frequently chosen than radiation for the majority of the genes (338 out of 466 genes with successful model training), corroborating a previous result with field data^9^. However, when the models were trained with GC or mixed data, radiation was more frequently used than temperature as the predictor of the majority of the genes (GC: 308/466 genes; mix: 378/466 genes). The choice of the predictor was also more consistent across data subsampling when we used GC or mixed data than field data (Fig. 3b).

**Fig. 3.**
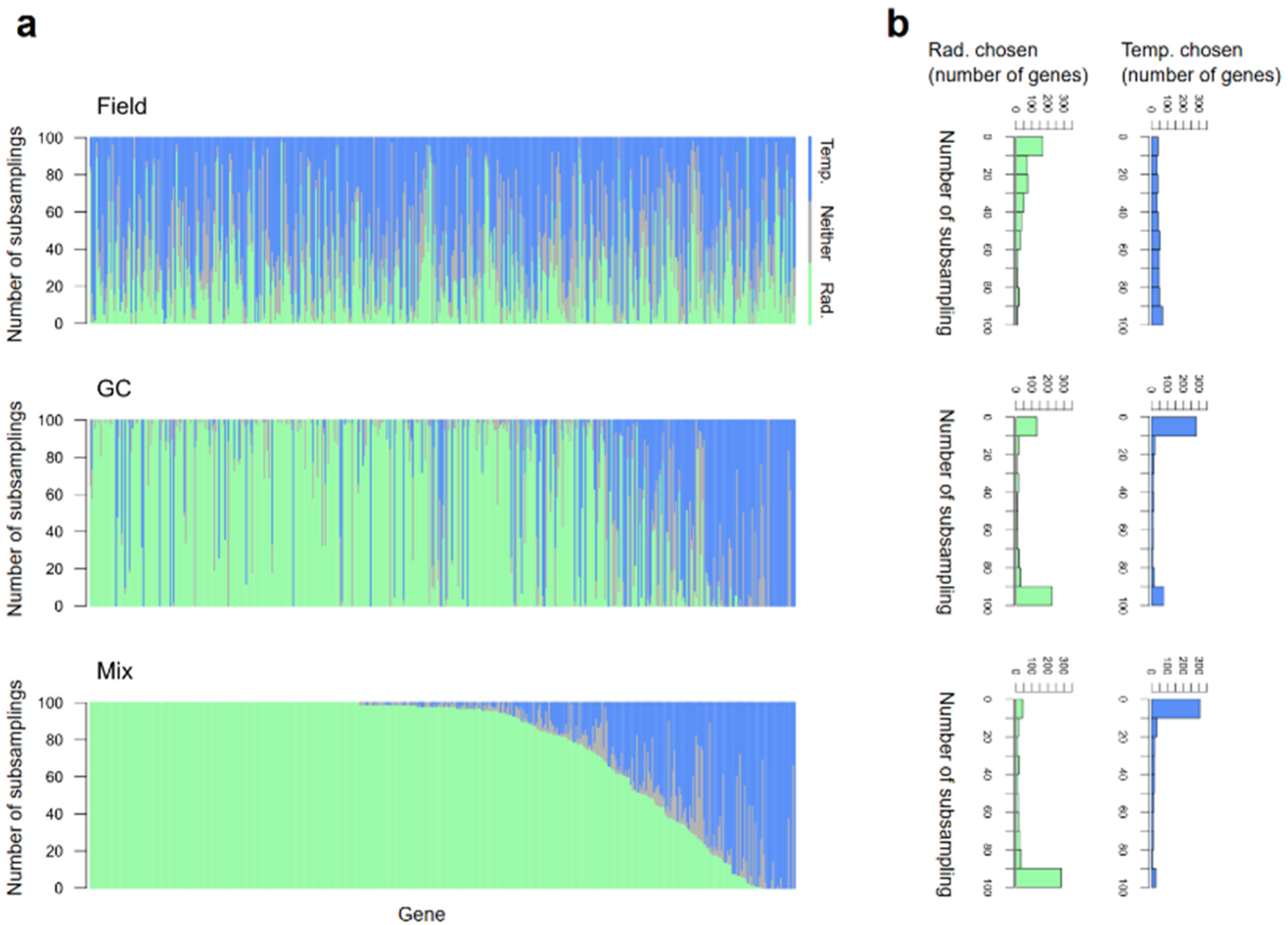
The number of times temperature or radiation (or neither) was chosen as the predictor of gene expression (Koshihikari). **a**, A single vertical line represents a gene whose green, blue, and grey parts show the frequency where radiation, temperature, or neither was chosen (among the 100 data subsampling). The 466 genes are sorted according to the frequency at which radiation was chosen in the model training with mixed (GC + field) data. **b**, Histograms showing how frequently temperature or radiation was chosen.

To identify the genes for which the input data influenced the choice of predictors, we extracted the representative genes for which temperature was consistently chosen with field-data training and radiation was consistently chosen with mixed data training (criterion for consistency: ≥ 80% subsampling). The number of representative genes extracted was 92 for Koshihikari and 101 for Takanari (Supplementary Table 8), where 38 genes were shared. Because individual genes represent clusters of different sizes (number of genes), we estimated the genome-wide impact; among the 16634 genes belonging to the 466 clusters, 3636 (21.9% of the total genes) for Koshihikari and 3988 genes (24.0%) for Takanari, with 1630 shared genes (9.8%).

### Mixed data improves the prediction of gene expressions in the field

To examine whether model predictions improve quantitatively due to the use of GC results in model training, we evaluated the models’ performances by applying the models to field-grown plants (*n* = 615) that were not included in the training data. Because the results can depend on training data sizes, we also varied the sizes of subsampled training data (subsampled 100 times for each size). We calculated mean absolute error (MAE) to measure the model performance by applying the trained models to the test data. Thus, we obtained 100 MAE values for 466 genes (Fig. 4a–c). Prediction performance varied substantially across genes, and these differences were relatively consistent across training data subsampling.

**Fig. 4.**
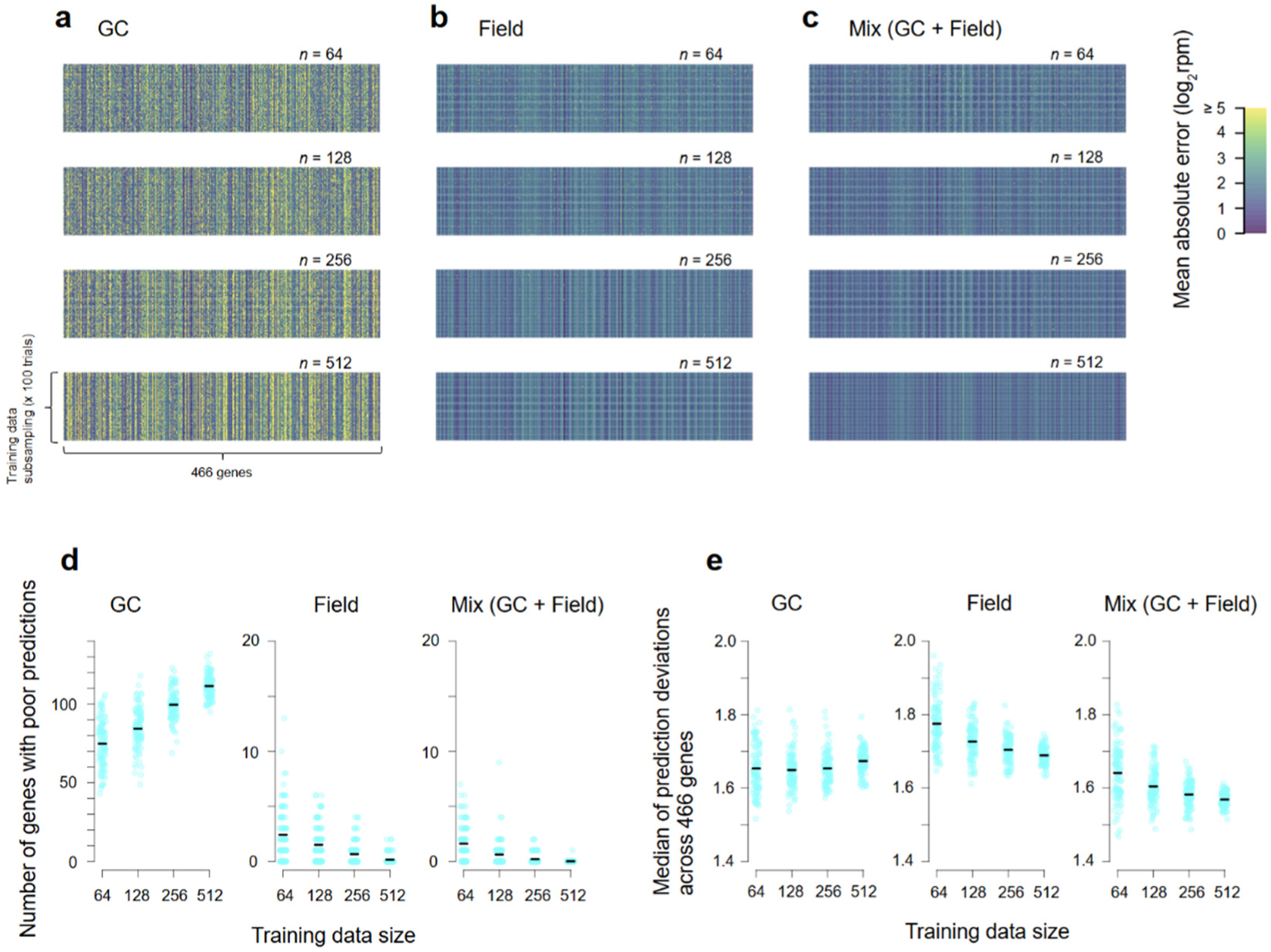
Prediction performances of gene expression models trained with different data sets. The performances were evaluated by applying the models to the field test data (not included in the training data). **a**–**c**, Heat maps of mean absolute errors (MAE). The GC + Field (c) models were trained with a mixture of GC and field data (50% each). **d**, The numbers of genes with poor predictions (log_2_rpm ≥ 5). **e**, Medians (across 466 genes) of the MAE values. In **d** and **e**, data points correspond to training data subsampling, and black horizontal lines represent means.

As a general trend, the use of mixed data (GC + Field) resulted in better predictions than other data sets (Fig. 4d). We considered MAE ≥ 5 (in log_2_rpm) as poor predictions. The number of genes with poor predictions almost always exceeded 50 when we used GC data alone for model training (Fig. 4d). The genes with poor predictions were much fewer with field-alone or mixed data. The number of genes with poor prediction decreased with increasing training data size for field and mixed training data. Notably, the largest mixed data resulted in the virtual absence of genes with poor predictions. Larger training data size for GC models decreased model performances, suggesting overfitting. Medians of the MAEs calculated across genes, which reflect the average performances of the models trained with different data sets, were small when the models were trained with large mixed data (Fig. 4e).

## Discussion

The correlation of irradiance and temperature in the field makes it difficult to evaluate the irradiance and temperature response of plants. In this study, by combining massive transcriptome analysis of rice leaves grown under controlled conditions and statistical modelling predicting rice transcriptome from meteorological data, we successfully distinguished the effect of irradiance and temperature on the plant transcriptome.

When we trained the models with field data, many gene-specific models chose temperature and radiation at intermediate frequencies (∼50), suggesting that the strong correlation between temperature and radiation in the field made it difficult to identify the environmental variable affecting the gene expression levels (Fig. 3). On the other hand, models trained with either the GC data or mixed data were more consistent in the choice of the environmental variables (∼0 or ∼100). These results suggest that using laboratory (i.e., GC) data where temperature and radiation do not correlate enables the models to choose the appropriate environmental variable to predict gene expression. However, model training only with the GC data led to poor predictions, with the evidence of overfitting with increasing training data size (Fig. 4). The GC data included only very young plants. In contrast, the field test data included plants of various ages. The models trained with the GC data thus extrapolated for plant age to make predictions. Field and GC data appear complementary to each other. GC data enabled the choice of better predictors (Fig. 3), but GC data alone led to poor parameter estimation (Fig. 4). The mixture of GC and field data has enabled the choice of better predictors and good parameter estimation, which explains the high prediction performance of the models trained with the mixed data. The test data we used were collected in a relatively similar environment to that of the field data for training. The performance differences between models trained with different data sets may become more pronounced when tested using data from novel environments (e.g., higher latitude).

Our results highlight the importance of irradiance in the transcriptome regulation. Previous studies using statistical modelling^9,10^ have suggested that temperature was more important than irradiance. A transcriptome study showed that the diurnal change of expression of *Brachypodium distachyon* was mainly affected by temperature^39^. In addition, an increase in the night temperature was found to have affected the diurnal change of transcriptome by regulation of PRR genes of rice panicles^40^. However, the temperature in these studies was confounded with irradiance, although continuous light/dark or temperature conditions exist in MacKinnon et al.^39^. This might have apparently increased the relative importance of temperature for transcriptome regulation in the previous studies.

Expression of some genes is known to be affected by both light and temperature^18,39,41^. On the other hand, our statistical model chose only one environmental parameter (irradiance or temperature or non) and cannot consider the joint regulation by both irradiance and temperature. Although it is possible to consider a model with multiple environmental factors, it would become more complex. The statistical model in Matsuzaki et al.^10^ considers both irradiance and temperature, but only 25 genes associated with the circadian clock were analyzed due to the high calculation cost. Using the model, Matsuzaki et al.^10^ showed that some circadian genes were affected by both irradiance and temperature, but more genes were affected by temperature than irradiance. This might be because field transcriptome and meteorological data, where irradiance and temperature correlate, were used for the model training.

The inclusion of the conditions where the temperature in the dark was higher than that in the light (Fig. 1a), sometimes called negative DIF (day-night temperature difference) may have contributed to improving the prediction of our model with mixed training data. Negative DIF conditions have been used to research plant circadian clock and phytohormone signalling^42,43^. In addition, negative DIF is also used to control the growth of horticultural crops^44^. However, few studies have evaluated the effect of the condition on plants utilizing the omics approach. Including negative DIF conditions in a sampling scheme seems effective in cutting off the correlation between temperature and irradiance, but it is unclear how much the negative DIF conditions contribute to improved prediction. Future studies will clarify the optimum experimental designs to improve the prediction of the model effectively.

Since the transcriptome data can be used for trait prediction^45–49^, improvement of the model used in this study will contribute to improving the accuracy of the trait prediction. Trait prediction is essential for the evaluation of the effect of climate change on crop production^50^. In this case, adding data obtained from extreme conditions (e.g., high or low temperature) will be effective. Improving trait prediction through the statistical model will contribute to understanding plant environmental response in the field and assessing climate change on plants.

## Methods

### Plant materials and growth conditions

In this study, we developed a high irradiance growth chamber (GC, ECP101, TECS Inc., Ibaraki, Japan) that can be controlled in parallel by one PC (Supplementary Fig. 1). The GC can control irradiance level and air temperature. The output value of each LED can be set from 0 to 100 %. In this study, we set the output value to 100%. The irradiance level was 1300 and 350 µmol photon m^-2^ s^-1^ (photosynthetic photon flux density [PPFD]) at the top (5 cm from the light source) and the bottom of the GC, respectively (Fig. 1a and Supplementary Fig. 1). The temperature was set to 15–40 °C. The GC can record temperature and relative humidity every minute.

A japonica rice (*Oryza sativa* L.) cultivar, ‘Koshihikari’, and an indica rice cultivar, ‘Takanari’, were used in this study. Seeds were sterilized in a 2.5% (v/v) sodium hypochlorite solution for 30 min and then soaked in water at 25 °C for 4 d. Germinated seeds were sown in a cell tray filled with nursery soil (N: P_2_O_5_:K_2_O = 0.6:1.2:1.0 g/kg). In the preculture, plants were grown for 14 days at 25/21°C and relative humidity of 55 ± 10 / 70 ± 10 % under a 12-h light/12-h darkness (12L/12D) photoperiod. Fluorescent lamps were used for the light source (FHF32EX-N-H, Iwasaki Electric Co., Ltd., Tokyo, Japan). The irradiance level was 400 ± 20 µmol photon m^-2^ s^-1^ (PPFD) at 5 cm from the light source (Supplementary Fig. 1). Plants were grown while maintaining plant height so that plant tips were 5 cm from the light source (Supplementary Fig. 1). Spot coolers were used to avoid burnt leaves due to the heat from the light source. Plants were then transferred to GC set to each condition (Fig. 1a). Environmental conditions in GC were composed of a combination of light period length (0L/24D, 8L/12D, 12L/12D, 16L/8D, and 24L/0D), light period temperature (20, 25, 30, 35, and 40°C) and dark period temperature (15, 20, 25, 30, and 35°C). Conditions with the same light and dark period temperature were excluded. As a result, experiments with 73 conditions were conducted (Fig. 1a and Supplementary Table 1). Experiments in 73 conditions were divided into 4 terms. In each term, sampling was conducted in 16, 23, 21, and 13 conditions (Supplementary Table 1). After transferring the plants to each condition, they were acclimatized for 48 h. Sampling was then conducted every 3 h for 24 h (1.5, 4.5, 7.5, 10.5, 13.5, 16.5, and 19.5h after the acclimation period, 8 times in total) (Fig. 1a). Under each condition, the uppermost, fully expanded leaves were sampled from one plant per sampling point, frozen in liquid nitrogen, and stored at −80°C for future use. At each sampling time point, sampling was completed within 20 min.

### RNA-Seq analysis

Frozen samples were homogenized with TissueLyser II (Qiagen, Hilden, Germany), and total RNA was extracted using the Maxwell 16 LEV Plant RNA Kit (Promega, Madison, WI, USA). RNA concentration was measured using the broad-range Quant-iT RNA Assay Kit (Thermo Fisher Scientific, Waltham, MA, USA). Total RNA (500 ng) was used as the input of each sample for library preparation. Library preparation for RNA sequencing was conducted using Lasy-Seq^51^ version 1.0 (https://sites.google.com/view/lasy-seq/) (Supplementary Figure S2). The library was sequenced using HiSeq 2500 (Illumina, San Diego, CA, USA) at Macrogen (Seoul, South Korea) with single-end sequencing lengths of 50 bp.

All obtained reads were trimmed using Trimmomatic version 0.36^52^ using the following parameters: TOPHRED33, ILLUMINACLIP: TruSeq3-SE.fa:2:30:10, LEADING:19, TRAILING:19, SLIDINGWINDOW:30:20, AVGQUAL:20, MINLEN:40, indicating that reads with more than 39 nucleotides and average quality scores over 19 were reported. Then, the trimmed reads were mapped onto the reference sequences of the IRGSP-1.0_transcript (2018-03-29)^53^, the mitochondria (NC_001320.1) and chloroplast (NC_011033.1) genomes, and the virus reference sequences, which were composed of complete genome sequences of 7457 viruses obtained from NCBI GenBank^6^ using RSEM version 1.2.21^54^ and Bowtie version 1.1.1^55^ with default parameters. The reads per million (rpm) were calculated using the nuclear-encoded gene raw count data, excluding the genes encoding rRNA, as described by Kashima et al.^6^. The number of reads was 0.18–6.91 million per sample (Supplementary Fig. 2a; Supplementary Table 2).

### Sample swap correction

We corrected sample annotations by identifying outliers in the temperature regression on the transcriptome (Supplementary Fig. 3). When we made a statistical model that predicted the temperature inside the GCs from the transcriptome, the result implied that some sample IDs were wrongly annotated, seemingly due to samples being swapped at 96-well plate level during library processing for RNA-Seq. To evaluate the accuracy of the temperature prediction, we split the data into those from a specific plate (test data) and the others (training data). Using the training data, we parameterized a generalized linear model (GLM) with Gaussian distribution that predicted the temperature inside the GCs from the transcriptome. We used the lasso (least absolute shrinkage and selection operator) technique for sparse regression. We adopted the regularization parameter *λ* that minimized MAE when fitted to the training data. By applying this parameterized GLM to the test data, we predicted the temperature (we used ‘glmnet’ package of R for this analysis). We repeated this procedure for all plates (Supplementary Fig. 3a). This analysis suggested that plates 9 and 11 (Supplementary Table 2) had been swapped during the lab work. It also indicated that plates 10 and 12 (Supplementary Table 2) had been similarly swapped. We re-annotated these samples, assuming that our inference was correct. We then repeated the same procedure (prediction of temperature). This second round of temperature prediction showed consistent, substantially improved accuracy and precision and did not imply a systematic sample interchange (Supplementary Fig. 3b). We thus assumed that the annotations used in this second prediction round were correct.

### Genotyping by sequence

We screened the genotypes of sequenced samples to correct possible sample swaps between cultivars (Supplementary Fig. 2b). Following the method of Kashima et al. ^6^, we genotyped samples by mRNA sequences. Briefly, we first compared sequences among samples and recoded the genotypes of each sample for all loci that were polymorphic in at least one of all possible sample pairs (247,645 loci). Among these loci, we used 26,746 loci known to show cultivar-specific SNP (a subset of previously established 1,427,878 loci with cultivar-specific SNP). Each locus of each sample was classified into one of four categories: Koshihikari genotype, Takanari genotype, unknown genotype (e.g. sequence error), or no sequence (number of sequenced loci were 4,005–20,953, with 12,786.8 on average). We calculated the proportions of loci with cultivar-specific genotypes among sequenced loci (Supplementary Fig. 2b). More than 70% of loci matched the nominal cultivar in all samples, but one, about 50% of loci showed genotypes different from both cultivars (among 13,634 sequenced loci). We discarded this sample from the main analyses.

### Statistical analysis of transcriptome data in 73 GC conditions

R software version 4.1.0 was used for the t-SNE, the calculation of amplitude of gene expression, DEG analysis, the calculation of Pearson’s correlation coefficient, co-expression gene network analysis, GO and KEGG enrichment tests^56^. A total of 17,742 genes in which the average log_2_(rpm+1) was > 1 were used for the analyses (Supplementary Fig. 2c). T-distributed Stochastic Neighbor Embedding (t-SNE) was conducted using the R package ‘Rtsne’ version 0.15^57^.

The diurnal oscillation of each gene expression (log_2_(rpm+1)) of each cultivar and condition were analyzed by the smooth.spline() of the R with the parameter spar set as 0.3 and intervals of 1.5 h. We defined the amplitude as the difference between the maximum and minimum values of the diurnal oscillation. A total of 12,770 genes whose average amplitudes of both Koshihikari and Takanari > 2 were used to compare the amplitudes between cultivar and conditions. Differentially expressed genes (DEGs) were extracted by paired t-test in which samples of Koshihikari and Takanari at the same time point are regarded as paired samples. FDR was controlled using Benjamini and Hochberg’s method^58^.

Co-expression gene network analysis was conducted by the R package ‘WGCNA’ version 1.70-3^35^. A signed network was constructed using the adjacency and TOMsimilarity function with power = 14. Modules were detected using the cutreeDynamic function, with deepsplit = 4 and minClusterSize = 20. Similar modules were then merged using the mergeCloseModules function with cutHeight = 0.2. As a result, 5889 genes were assigned to 22 modules. Module Eigengene (ME), the first component in PCA of the gene expression profiles, was calculated for each module. The constructed networks were visualized using Cytoscape version 3.9.0^59^. Genes with adjacency > 0.05 are shown. To explore the hub genes that can have a central role in the gene co-expression network regulation in module 5, we used NetworkAnalyzer in Cytoscape. Three genes with Degree > 4 and Betweenness Centrality > 0.3 were extracted as hub genes.

Gene enrichment tests for GO and Kyoto Encyclopedia of Genes and Genomes (KEGG)^60^ pathways were conducted using the R package ‘GO.db’ version 3.13.0^61^ and ‘KEGGREST’ version 1.32.0^62^, respectively, as described by Nagano et al.^7^. The FDR was controlled using Benjamini and Hochberg’s method^58^ with FDR = 0.05.

### Selection of representative genes

We used microarray data of rice leaves sampled in paddy fields in 2008^9^; we performed affinity propagation clustering using 17616 genes with expression greater than 2^5^ in more than 80% of the samples. 500 clusters were obtained from the affinity propagation clustering using the R package ‘apcluster’^63^. The exemplars of each cluster are considered as representative genes.

Since the gene annotation used for the RNA-Seq in this study^53^ was updated from those used in the microarray of Nagano et al.^9^, we updated the gene ID of the genes based on the homology search using BLASTn for nucleotides^64^. Among the 17616 genes, 16634 genes exist in the gene annotation used for the RNA-Seq; 24 among 500 representative genes obsoleted. We used the remaining 474 genes for the following analysis.

### Modelling gene expression dynamics

We used the R package ‘FIT’^8^, which trains a model of gene expression dynamics as a function of meteorological time series, time of day, and plant age. Gene expression dynamics were modelled independently for the two cultivars, Koshihikari and Takanari. We trained the models using the GC data (Koshihikari: *n* = 584; Takanari: *n* = 583), field data (Koshihikari: *n* = 805; Takanari: *n* = 714), or a mixture of them (50% GC and 50% field). When model training failed in any data subsampling with any training data, such genes were excluded from subsequent analyses (genes were omitted independently between cultivars). The prediction performances of the models were tested using field data that was not included in the training data (615 data points for Koshihikari and 539 points for Takanari). We allowed the models to use the meteorological data up to 72 hours ago.

## Data availability

The scripts used in this study are available at https://github.com/naganolab/GC_73_conditions. All datasets generated and/or used in this study are available in the Sequence Read Archive (SRA) under accession number PRJNA1074001.

## Acknowledgements

We thank Tex Inc. (Ibaraki, Japan) for the establishment of the GC, Fumie Kobayashi and Kyoko Mogami for technical assistance with the RNA-Seq experiments, and Dynacom Co., Ltd. (Chiba, Japan) for technical assistance with the analysis of RNA-Seq data. This study was supported by JST CREST Grant Number JPMJCR15O2, JST FOREST Grant Number JPMJFR210B, JSPS (JP20H00423, JP23H00386, and JP23K18156) and MEXT (JP23H04967) awarded to AJN.

## Author contributions

These authors contributed equally: YH. and DK. AJN, YH, and DK conceived the study. TT developed the GC. YH, SET, NM, and HW conducted the GC experiments. SET conducted the RNA-Seq experiments. YH and DK analyzed the data. YH, DK, and AJN wrote the manuscript with input from all co-authors.

## Supplementary Information

**Supplementary Fig. 1** GC and the preculture condition.

**Supplementary Fig. 2** Workflow of the RNA-Seq data preprocessing.

**Supplementary Fig. 3** Swap correction.

**Supplementary Fig. 4** Normalized expressions of eigengenes of each module.

**Supplementary Fig. 5** Expression levels of a gene encoding *Hsp 70* (*Os01g0840100*).

**Supplementary Fig. 6** The number of times temperature or radiation (or neither) was chosen as the predictor of gene expression (Takanari).

**Supplementary Fig. 7** Prediction performances of gene expression models trained with different data sets in Takanari.

**Supplementary Table 1.** 73 environmental conditions used in this study.

**Supplementary Table 2.** Sample attributes used in this study.

**Supplementary Table 3.** Enriched gene ontology in each module.

**Supplementary Table 4.** Enriched KEGG pathway in each module.

**Supplementary Table 5.** List of genes in module 5.

**Supplementary Table 6.** List of genes with positive correlation in expression with air temperature.

**Supplementary Table 7.** List of genes with negative correlation in expression with air temperature.

**Supplementary Table 8.** List of the representative genes whose predictors were temperature in the models trained with field data but radiation in more than 80% of the models trained with mixed data.

