## Supplementary Figure 1 for "Field-crop transcriptome models are enhanced by measurements in systematically controlled environments"

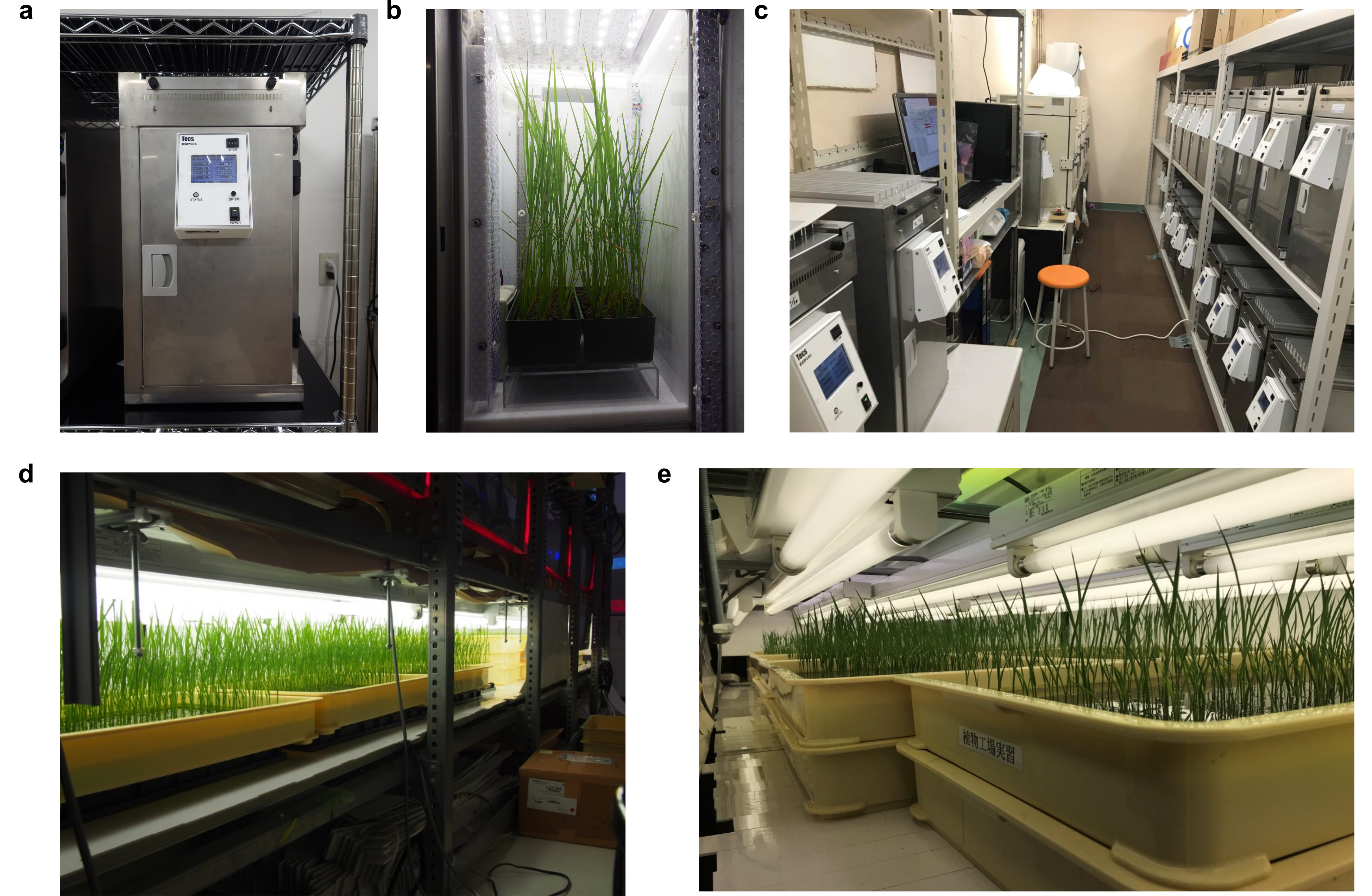
**Supplementary Fig. 1 GC and the preculture condition. a**, Front view of GC. b, Rice grown in GC. **c**, The parallel control of GC. **d**, Overview of the preculture condition. **e**, Rice grown in the preculture condition.

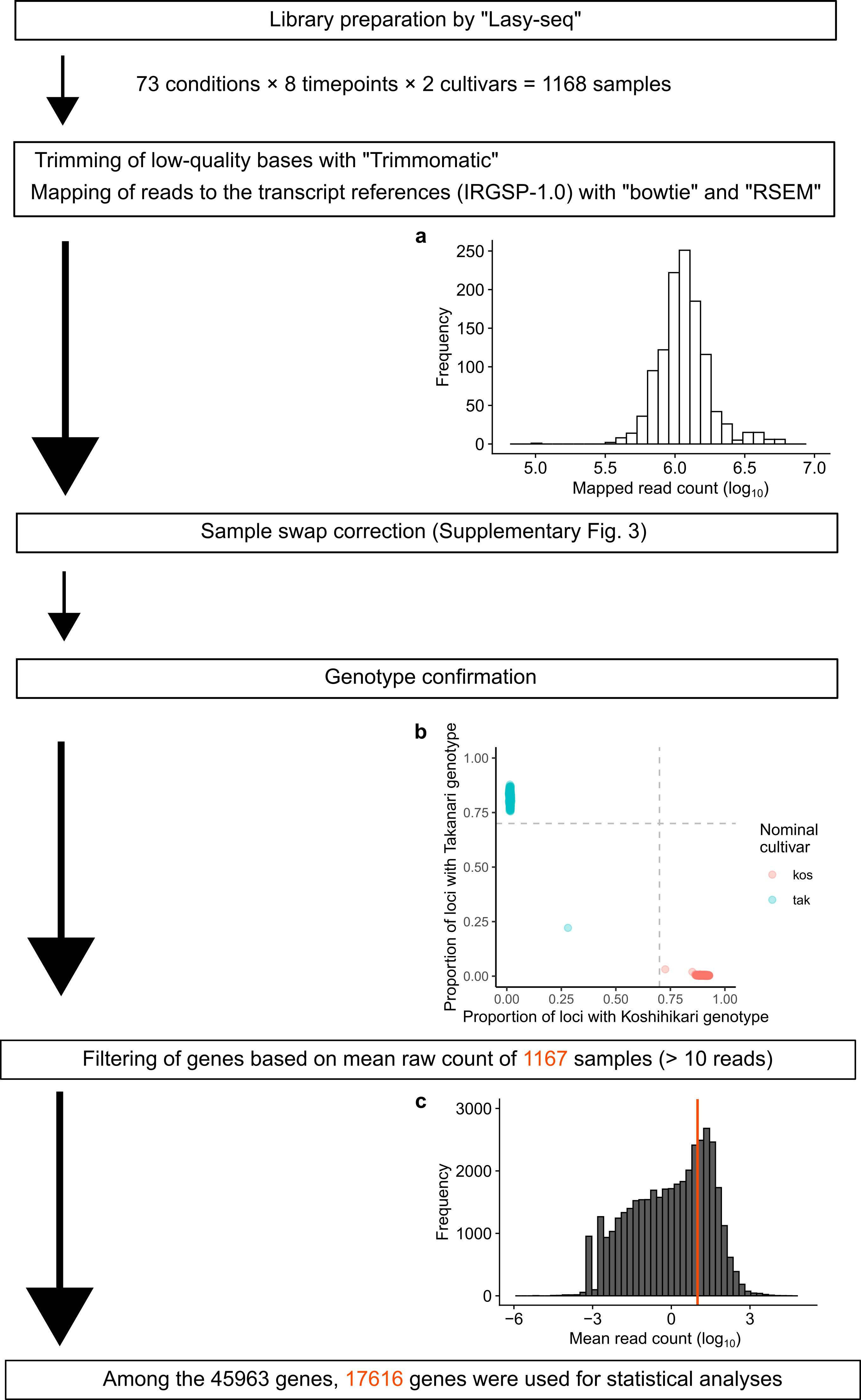

**Supplementary Fig. 2 Workflow of the RNA-Seq data preprocessing.** **a**, Histogram of the mapped read counts used for the calculation of rpm for each sample. **b**, Genotyping by sequence. Position of each sample in the two-dimensional genotype space. One sample was omitted based on the genotyping. **c**, Histogram of the mean read count for each gene. After filtering, 17,616 genes were used for the analyses.

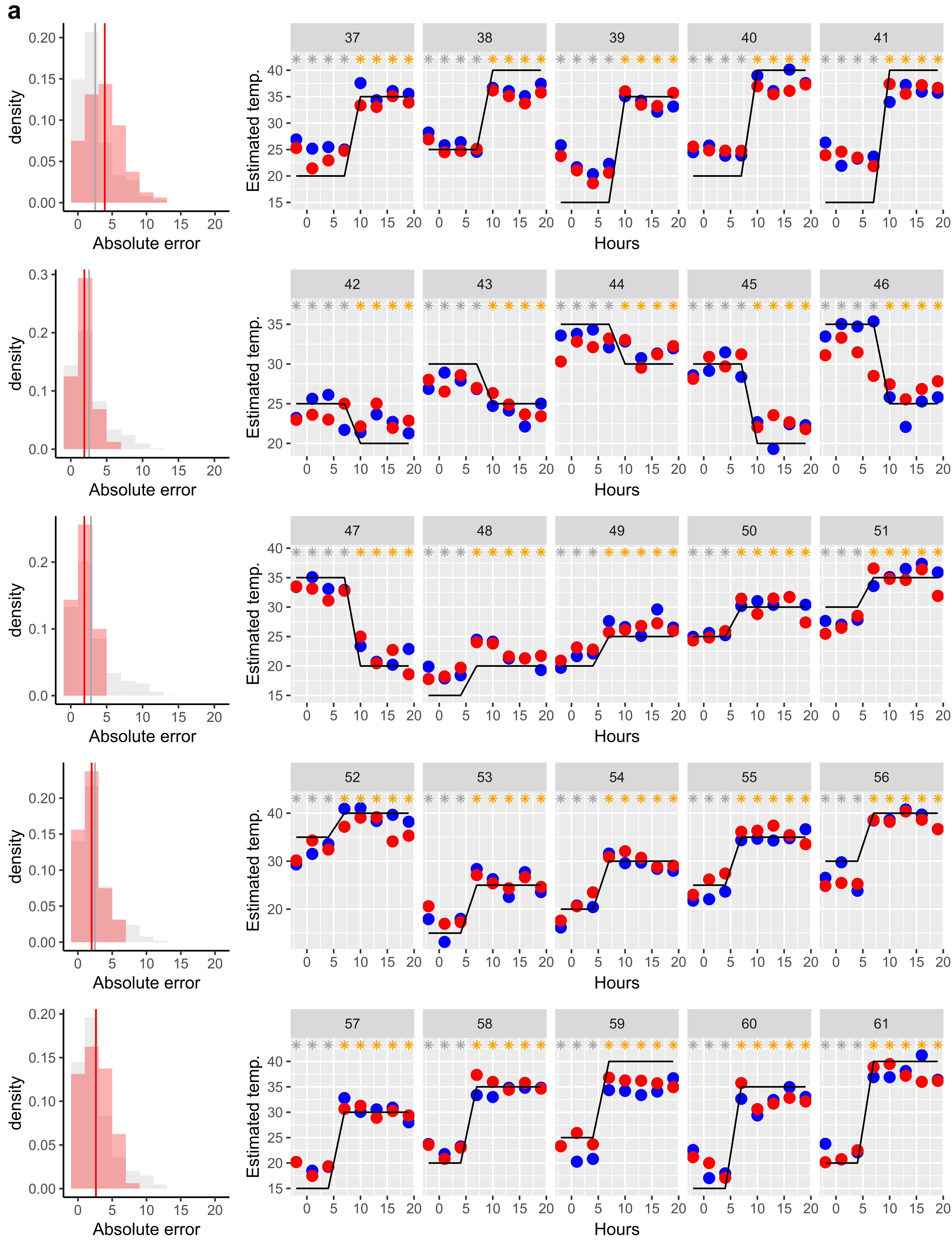

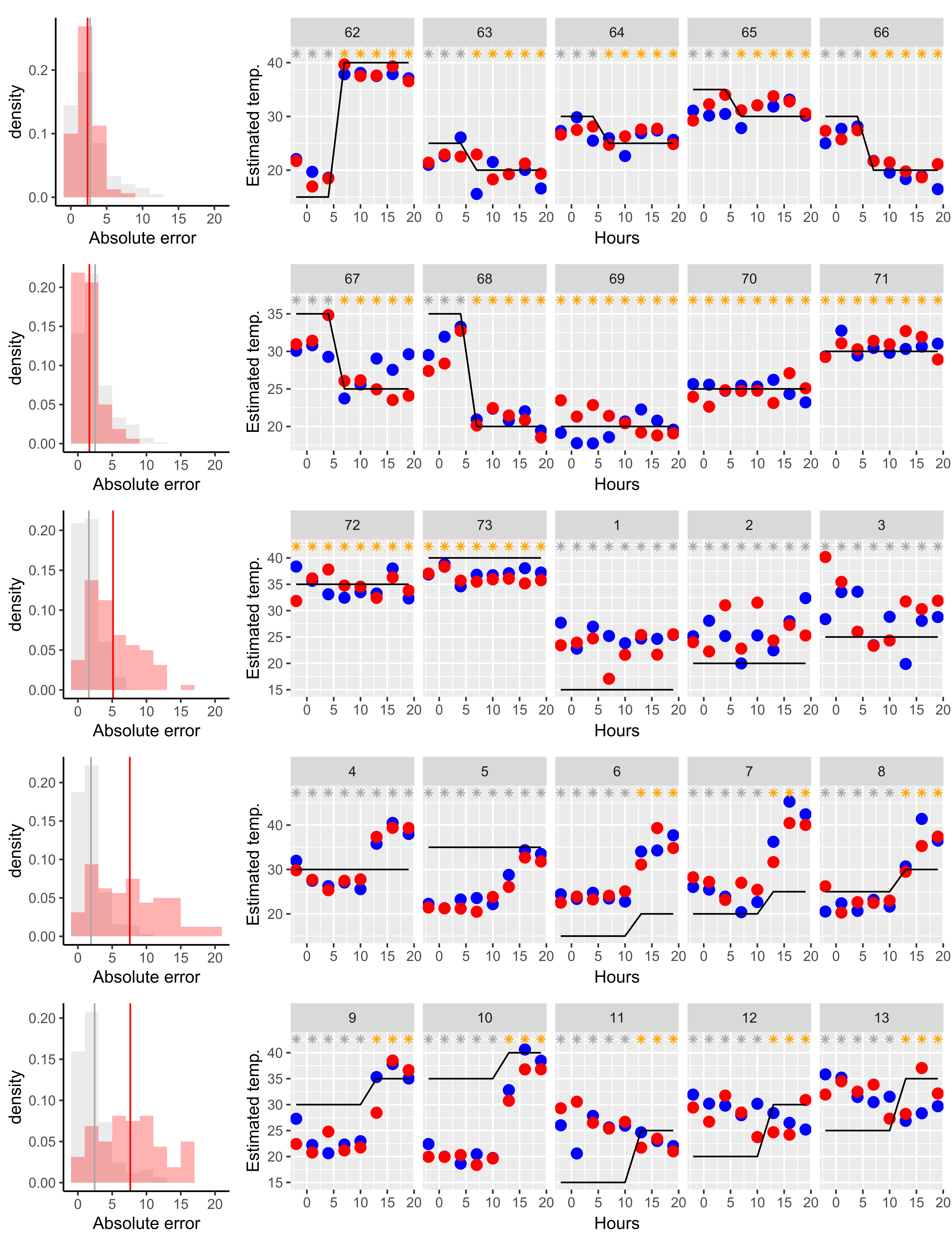

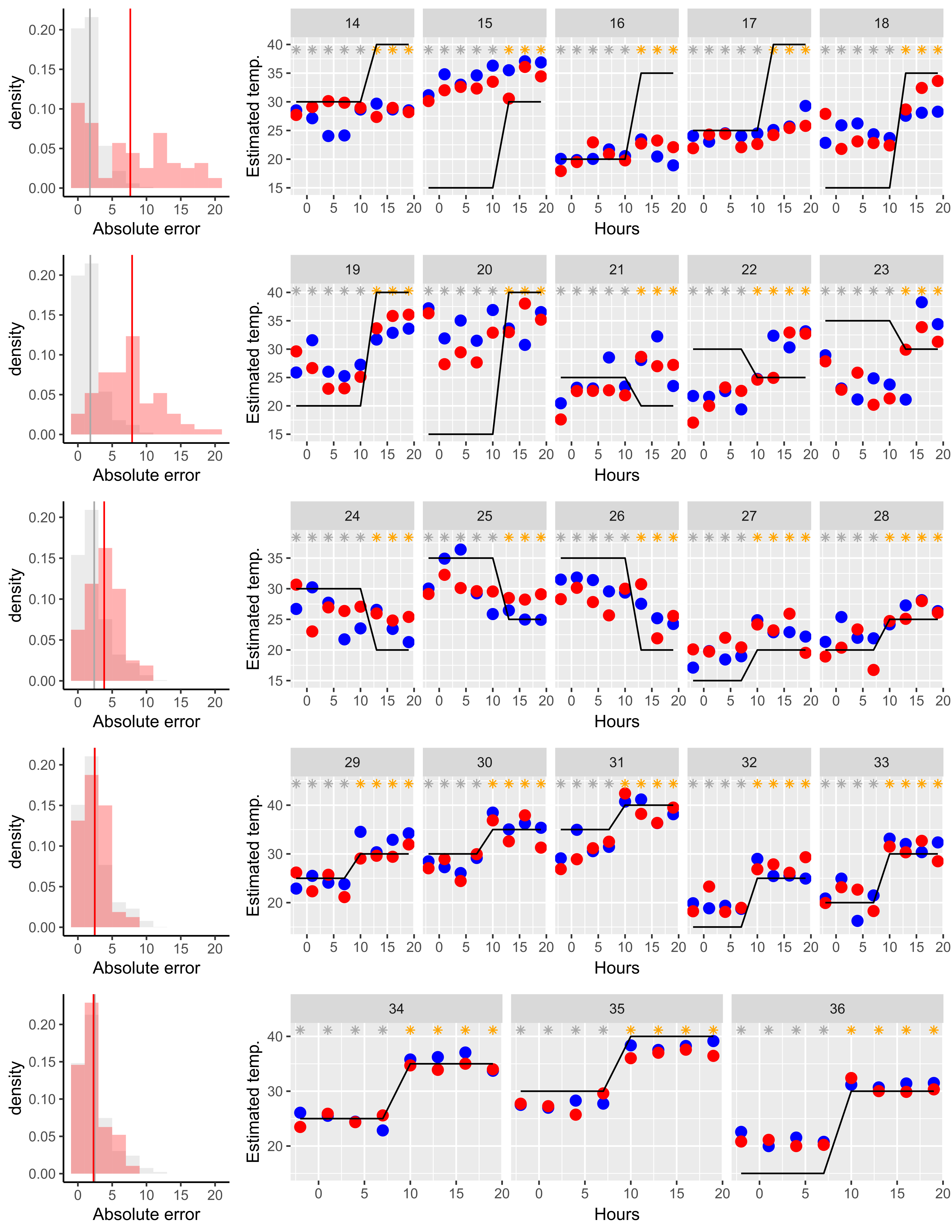

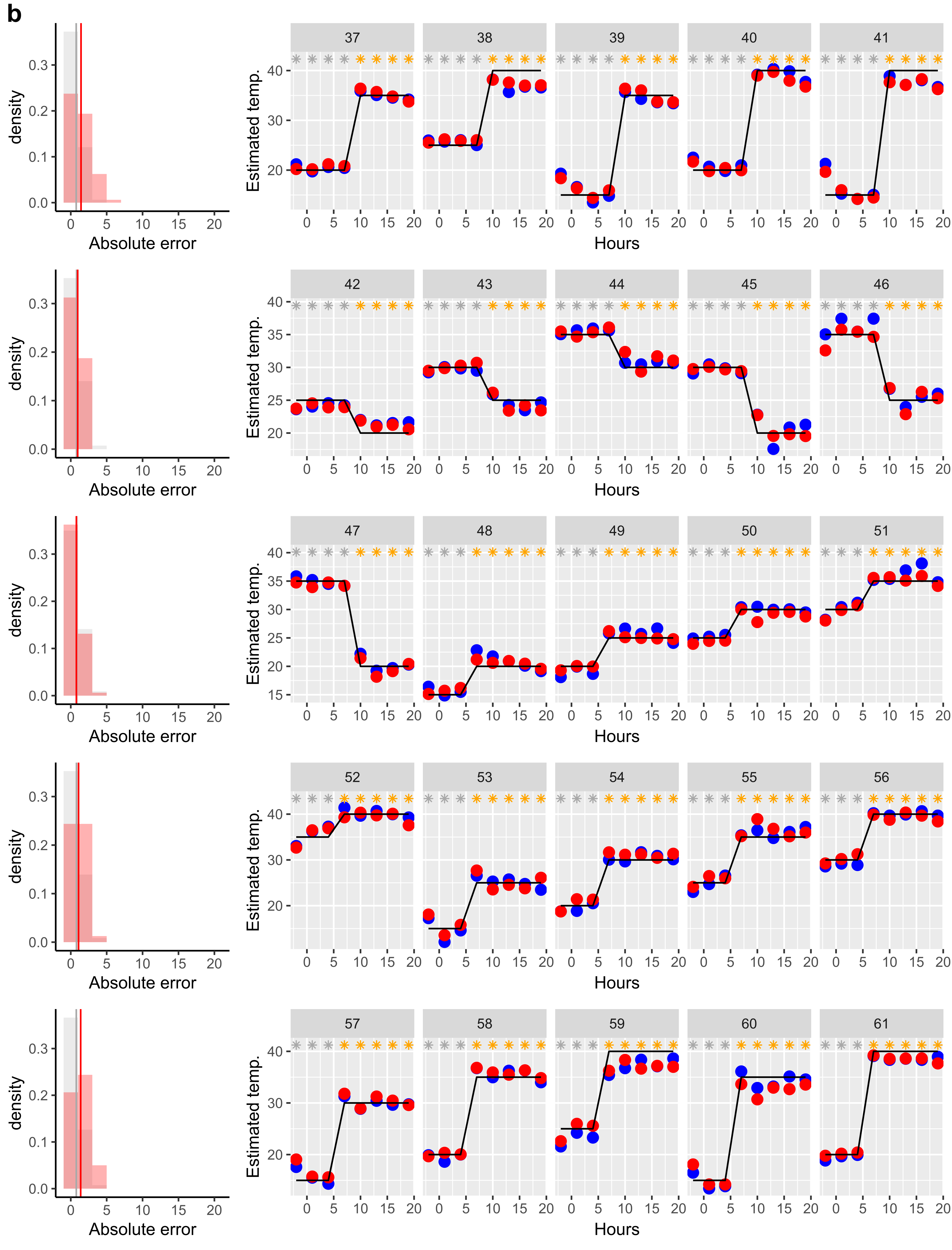

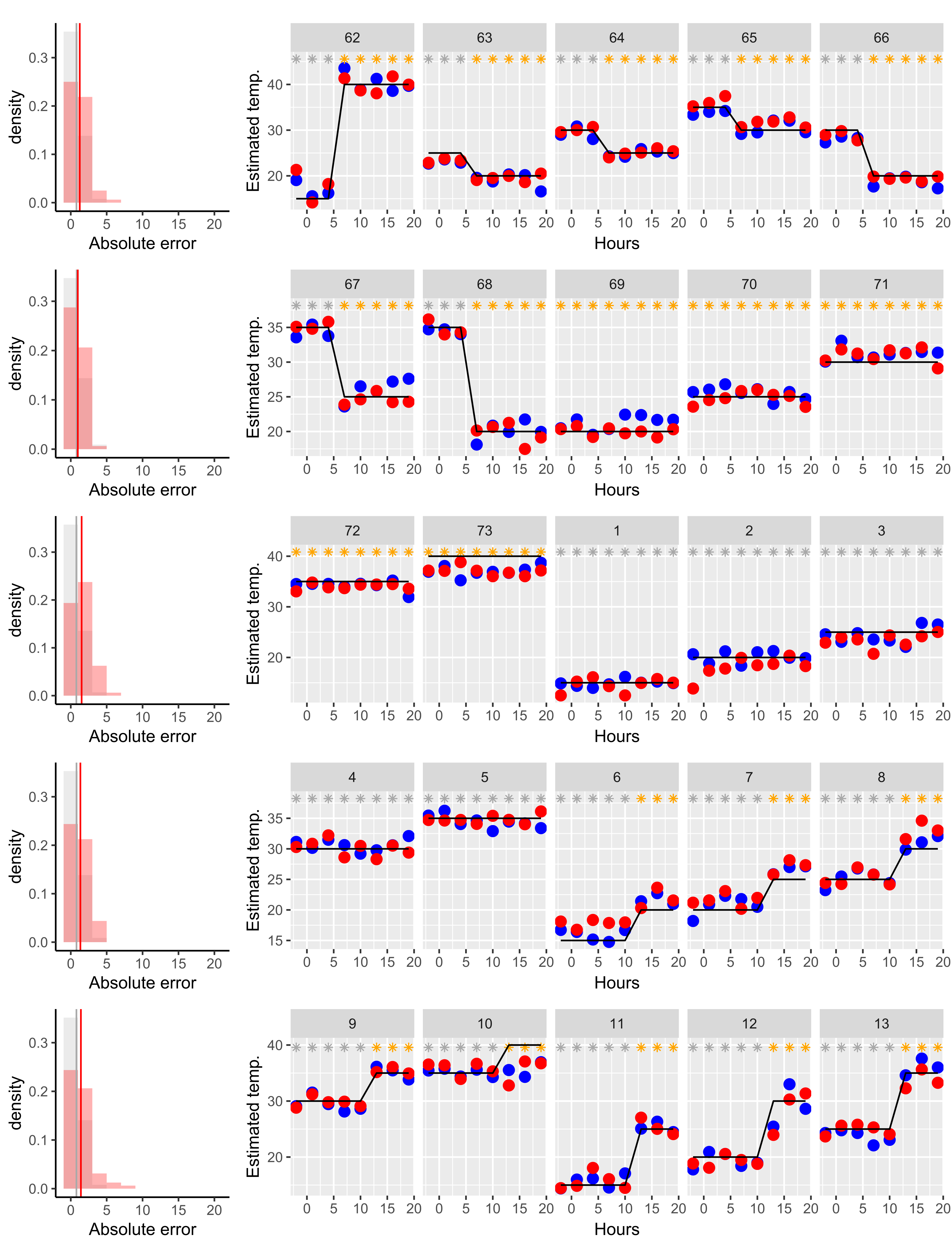

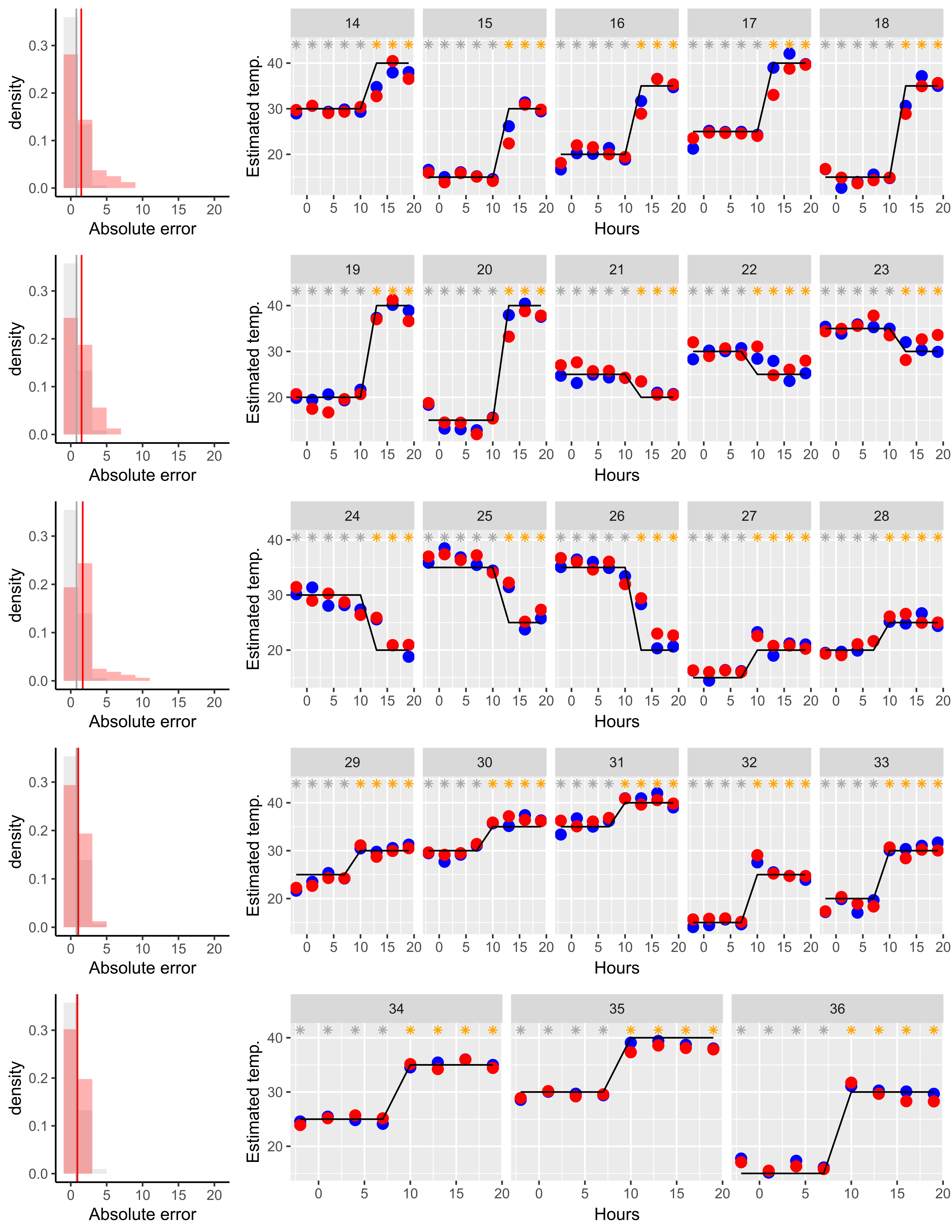

**Supplementary Fig. 3 Swap correction.** Performance of statistical models predicting the temperature inside the GCs during the first (**a**) and second (**b**) rounds. Each panel corresponds to a 96-well plate. In each panel, time series plots represent the model predictions (blue: Koshihikari, red: Takanari), and solid lines represent GC settings. Grey and orange asterisks represent light-dark cycles (grey: dark, orange: light). The numbers on the top of time series plots are experimental batches. Histograms show the absolute error distributions (red: that for the test data, grey: that for the training data). Vertical lines on the histograms represent mean absolute errors.

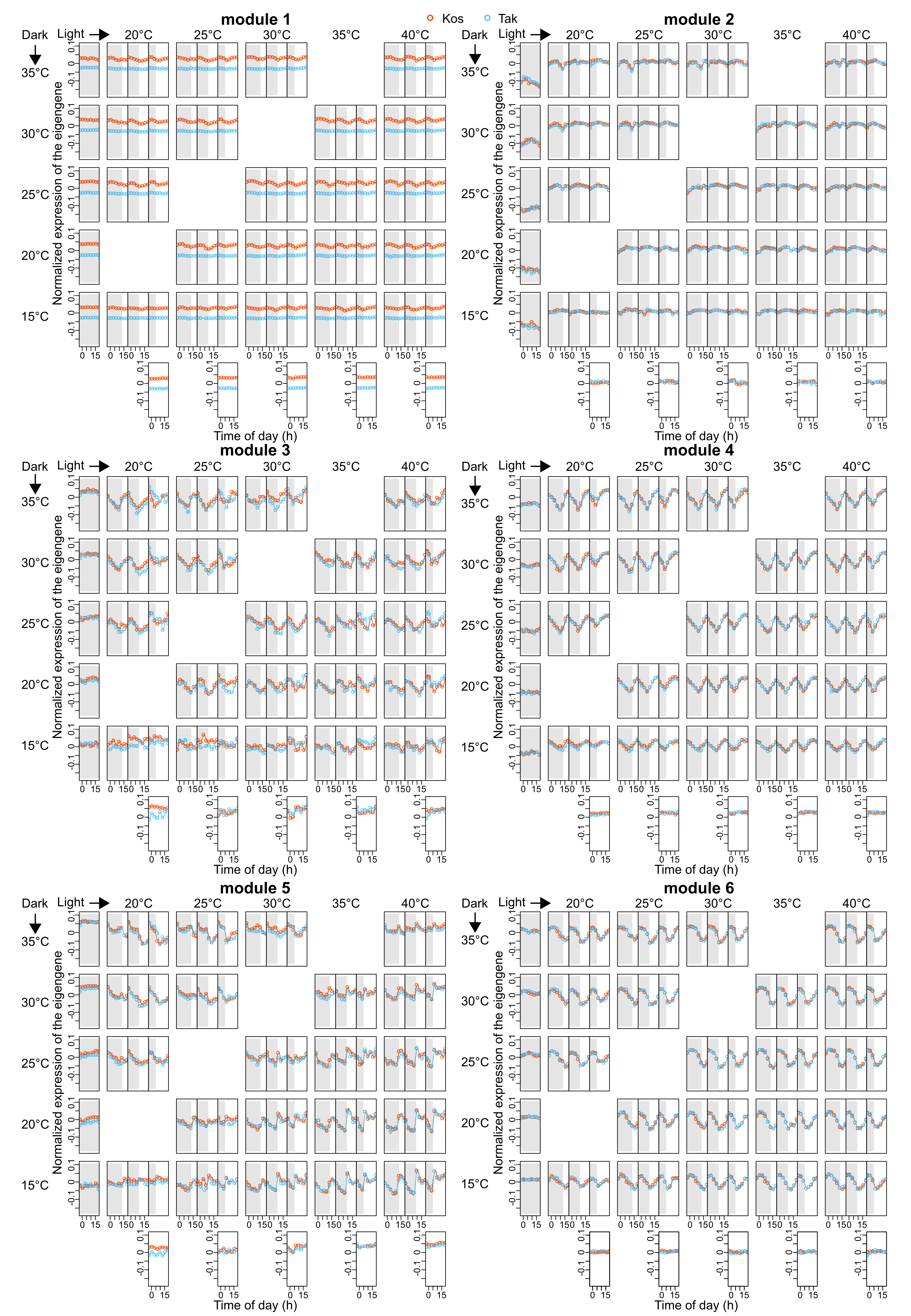

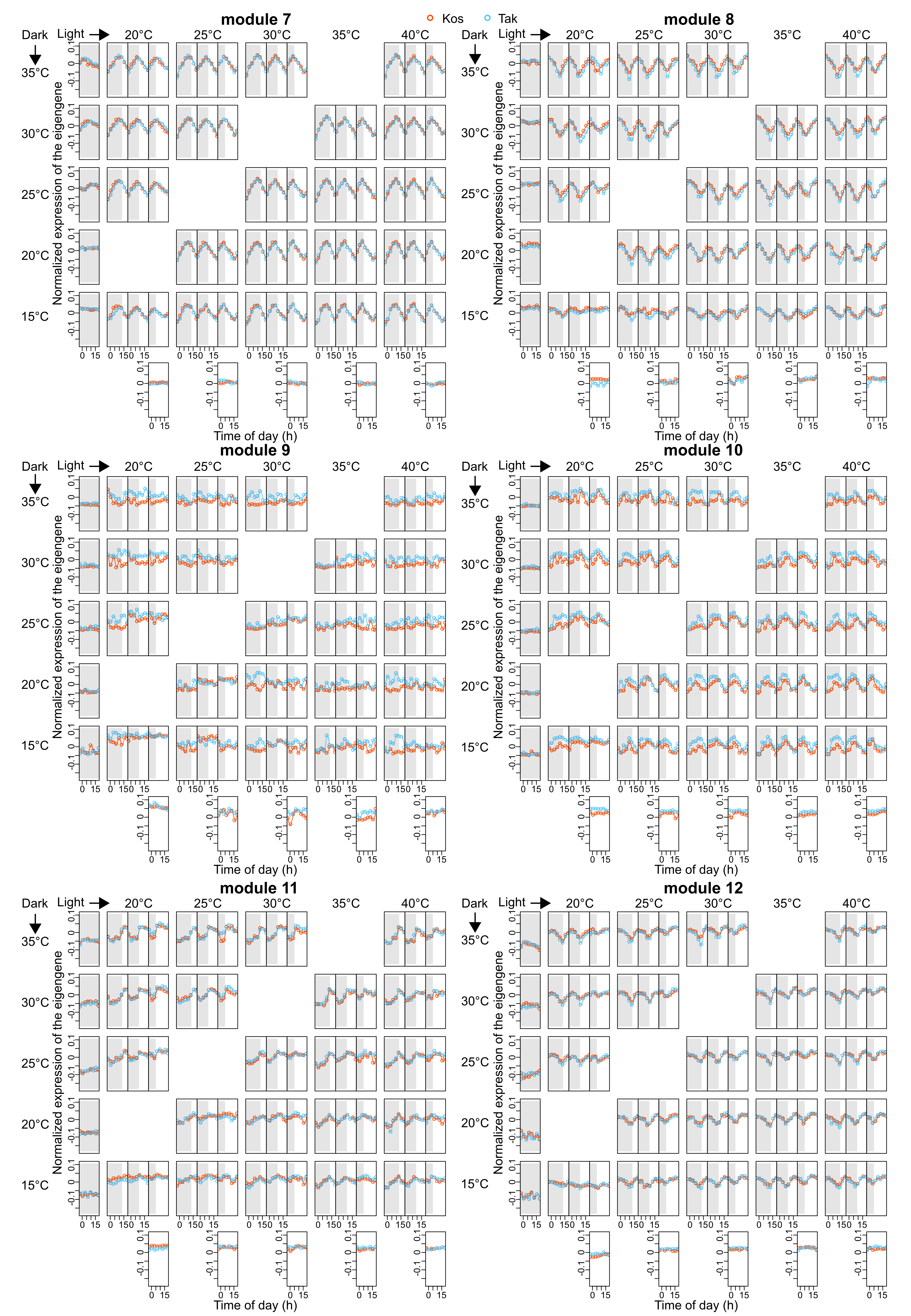

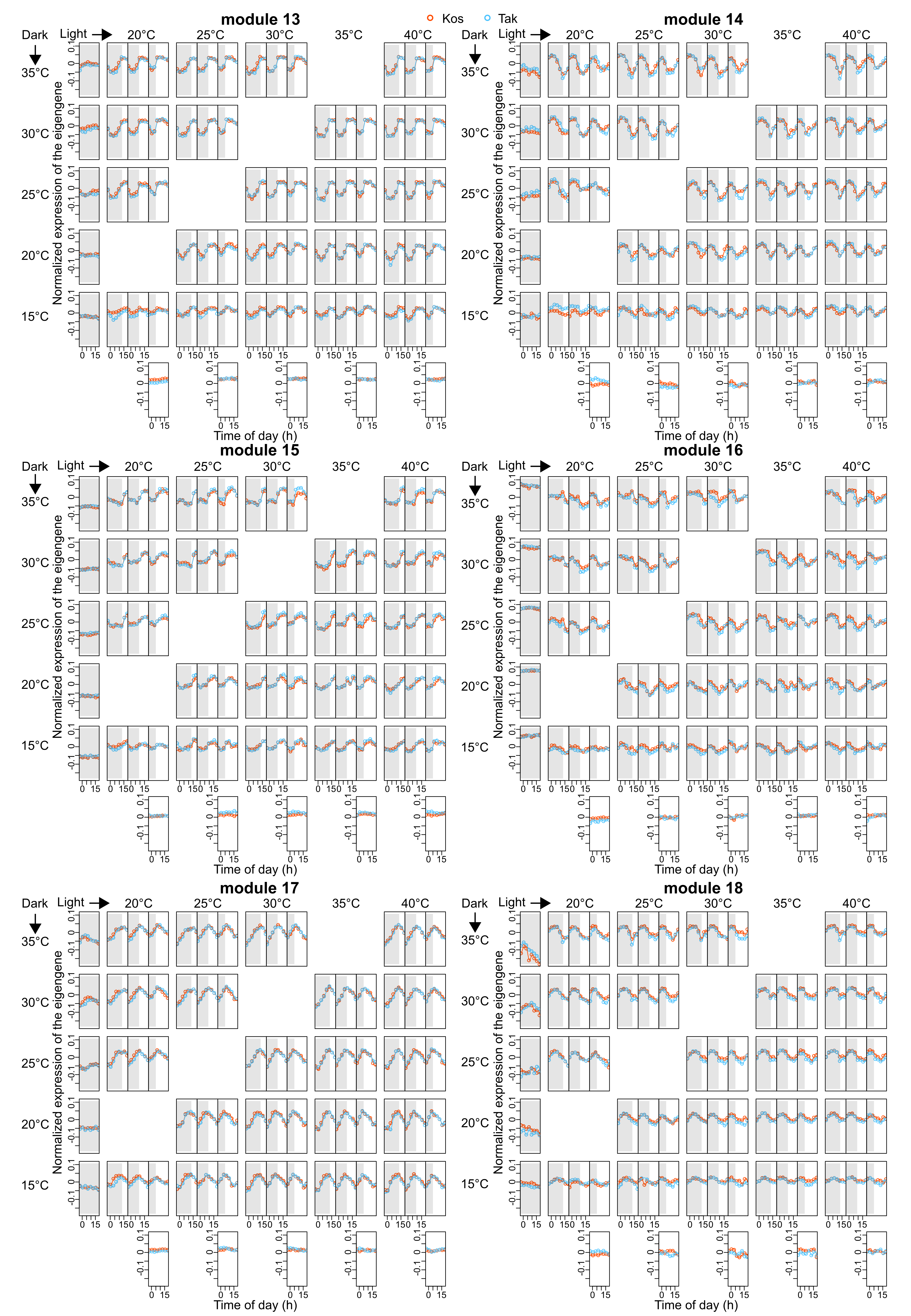

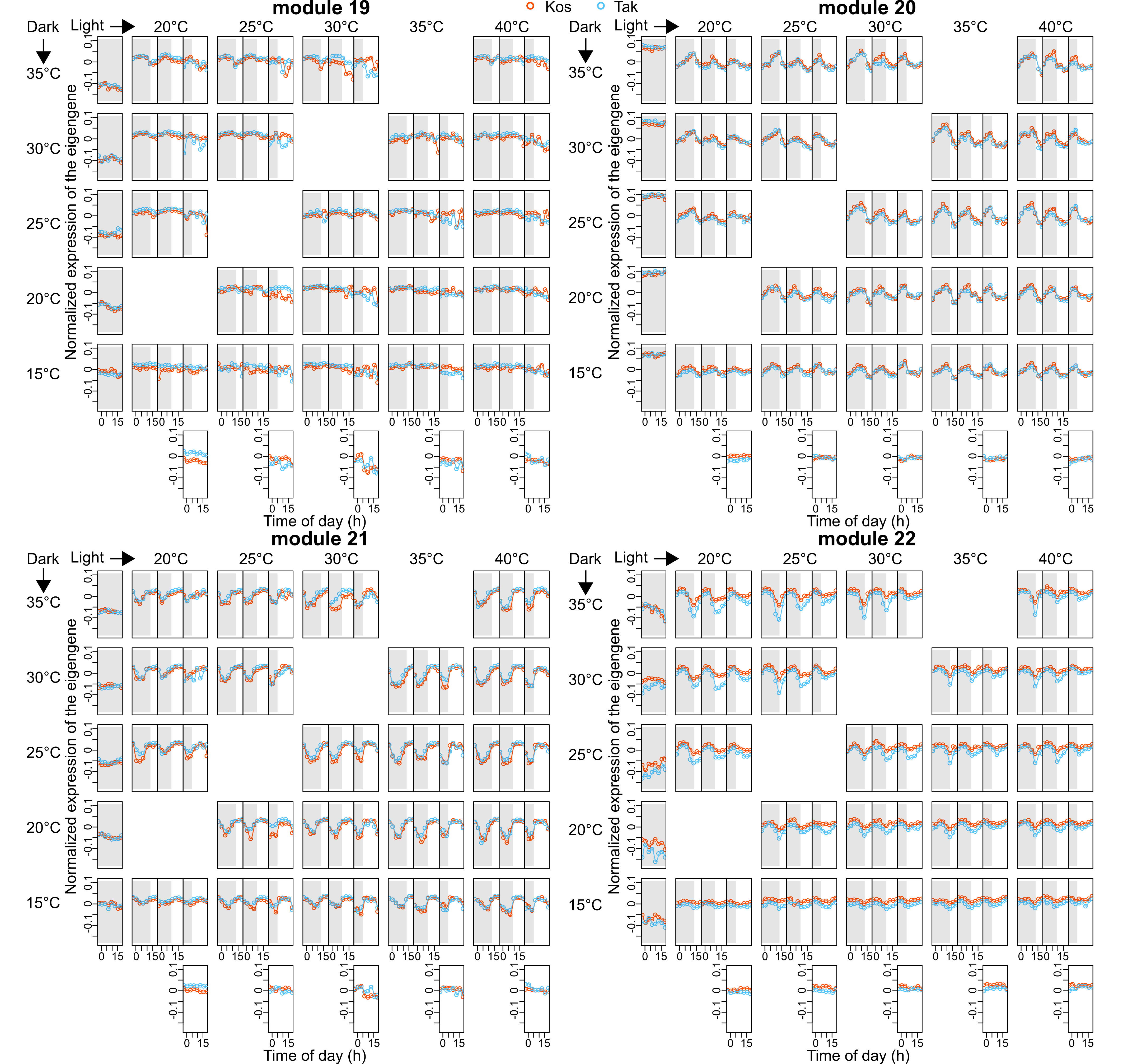

**Supplementary Fig. 4 Normalized expressions of eigengenes of each module.** The expressions of eigengenes were normalized using z-scores.

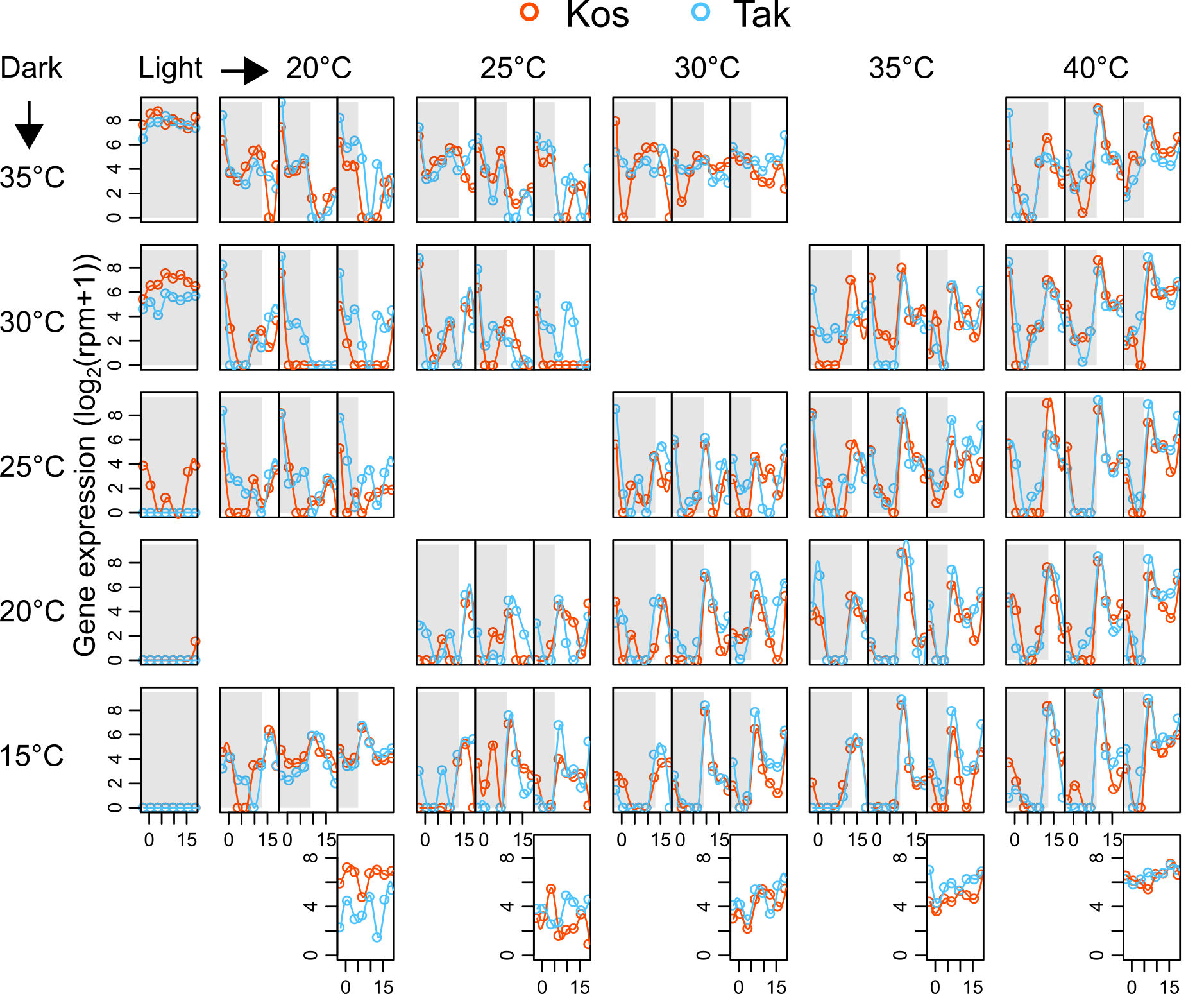

**Supplementary Fig. 5 Expression levels of a gene encoding *Hsp 70* (*Os01g0840100*).**

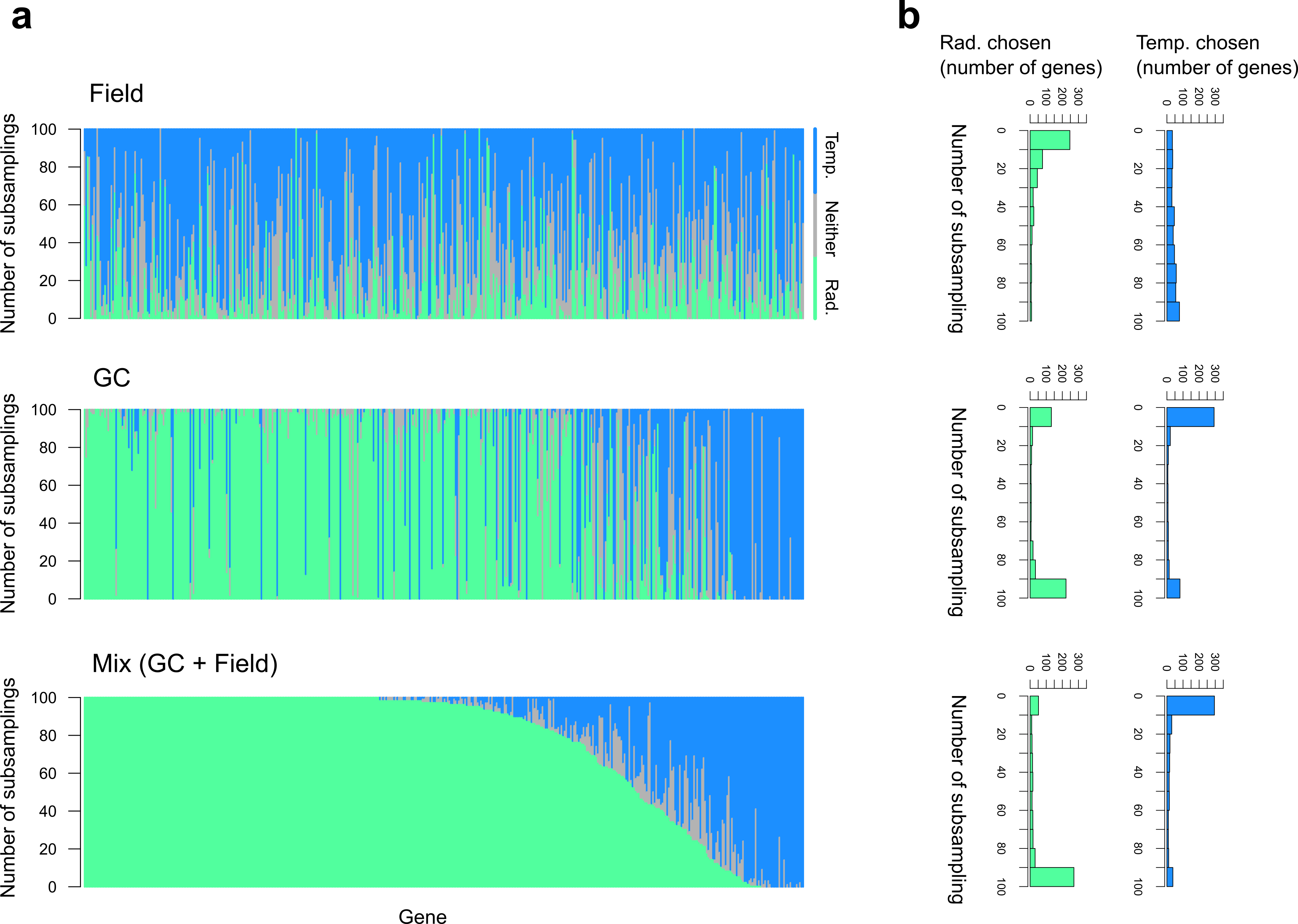

**Supplementary Fig. 6 The number of times temperature or radiation (or neither) was chosen as the predictor of gene expression (Takanari).** **a**, A single vertical line represents a gene whose green, blue, and grey parts show the frequency where radiation, temperature, or neither was chosen (among the 100 data subsampling). The 466 genes are sorted according to the frequency at which radiation was chosen in the model training with mixed (GC + field) data. **b**, Histograms showing how frequently temperature or radiation was chosen.

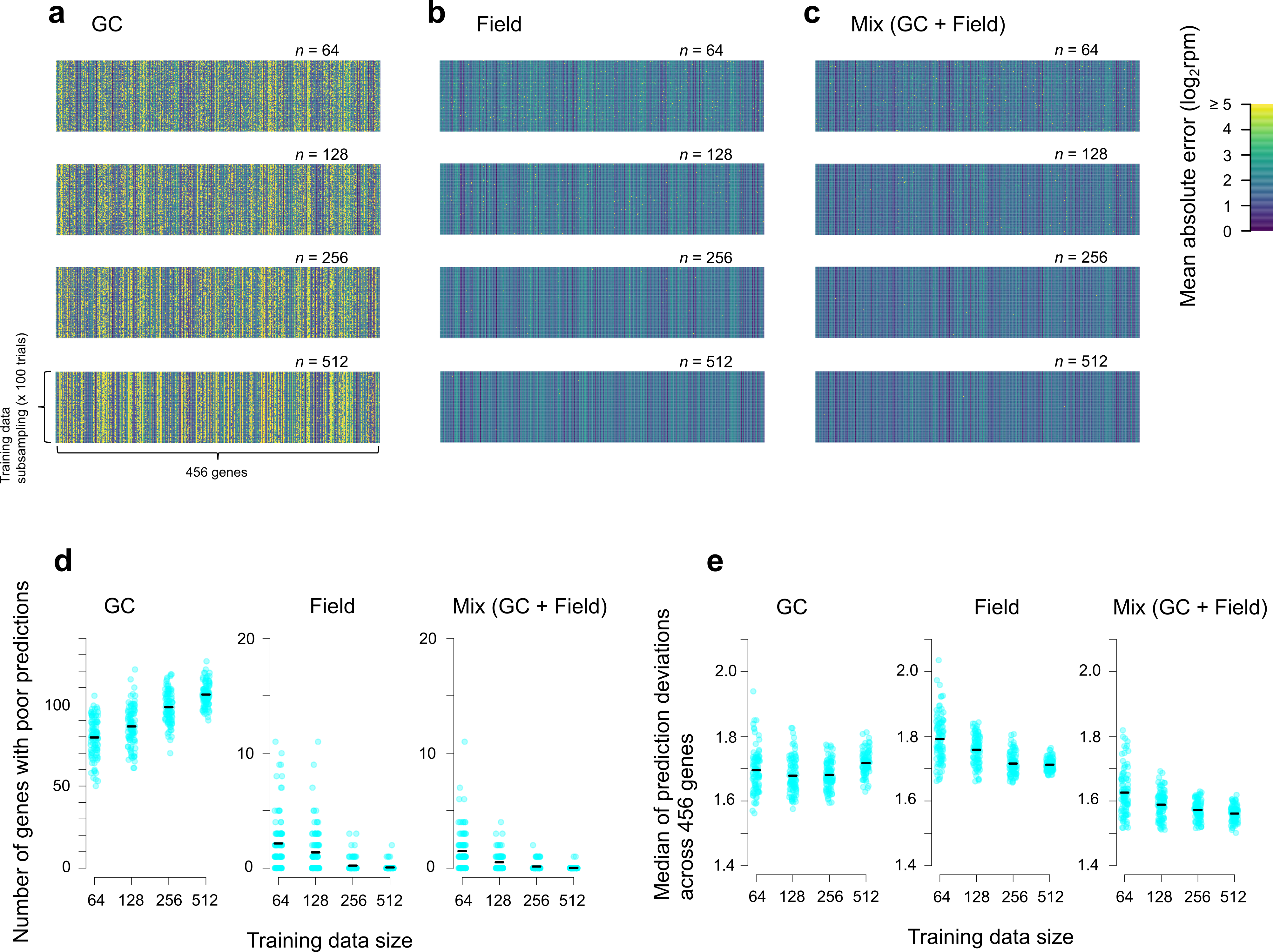

**Supplementary Fig. 7 Prediction performances of gene expression models trained with different data sets in Takanari.** The performances were evaluated by applying the models to the field test data (not included in the training data). **a**–**c**, Heat maps of mean absolute errors (MAE). The GC + Field (c) models were trained with a mixture of GC and field data (50% each). **d**, The numbers of genes with poor predictions (log2rpm ≥ 5). **e**, Medians (across 466 genes) of the MAE values. In **d** and **e**, data points correspond to training data subsampling, and black horizontal lines represent means.
